# From local spectral species to global spectral communities: a benchmark for ecosystem diversity estimate by remote sensing

**DOI:** 10.1101/2020.11.04.367995

**Authors:** Duccio Rocchini, Nicole Salvatori, Carl Beierkuhnlein, Alessandro Chiarucci, Florian de Boissieu, Michael Förster, Carol X. Garzon-Lopez, Thomas W. Gillespie, Heidi C. Hauffe, Kate S. He, Birgit Kleinschmit, Jonathan Lenoir, Marco Malavasi, Vítězslav Moudrý, Harini Nagendra, Davnah Payne, Petra Šímová, Michele Torresani, Martin Wegmann, Jean-Baptiste Féret

## Abstract

In the light of unprecedented change in global biodiversity, real-time and accurate ecosystem and biodiversity assessments are becoming increasingly essential. Nevertheless, estimation of biodiversity using ecological field data can be difficult for several reasons. For instance, for very large areas, it is challenging to collect data that provide reliable information. Some of these restrictions in Earth observation can be avoided through the use of remote sensing approaches. Various studies have estimated biodiversity on the basis of the Spectral Variation Hypothesis (SVH). According to this hypothesis, spectral heterogeneity over the different pixel units of a spatial grid reflects a higher niche heterogeneity, allowing more organisms to coexist. Recently, the spectral species concept has been derived, following the consideration that spectral heterogeneity at a landscape scale corresponds to a combination of subspaces sharing a similar spectral signature. With the use of high resolution remote sensing data, on a local scale, these subspaces can be identified as separate spectral entities, the so called “spectral species”. Our approach extends this concept over wide spatial extents and to a higher level of biological organization. We applied this method to MODIS imagery data across Europe. Obviously, in this case, a spectral species identified by MODIS is not associated to a single plant species in the field but rather to a species assemblage, habitat, or ecosystem. Based on such spectral information, we propose a straightforward method to derive *α*- (local relative abundance and richness of spectral species) and *β*-diversity (turnover of spectral species) maps over wide geographical areas.

## 1 Introduction

### 1.1 A quest for robust and reproducible *α*- and *β*-diversity measurement

The variability of life on Earth is heterogeneously distributed across the planet; in fact, ecologists and biogeographers have long questioned the potential causes of biodiversity distribution. Recently, the speed of change and the uncertainty about possible consequences on biodiversity is concerning to the global scientific community. The perception of these processes translates into the need to use standardized methods for biodiversity assessment and monitoring in order to gain a better understanding and identify general trends.

There is open debate as to the most reliable metrics for assessing biodiversity (see Jurasinski et al. (2009); Tuomisto (2010)). Until now, no consistent definition exists. Even the definition according to the Convention on Biological Diversity (CBD, 1992, https://www.cbd.int/convention/text/) is more confusing than clear: “Biological Diversity means the variability among living organisms from all sources, including, inter alia, terrestrial, marine and other aquatic ecosystems and the ecological complexes of which they are part; this includes diversity within species, between species and of ecosystems.” Biodiversity obviously includes quantitative (number of species, alpha-diversity, gamma-diversity), qualitative (turnover, composition, beta-diversity) and functional (complexity, trophic levels, ecosystem services) aspects. To sum up our understanding on the term biodiversity (i.e. biological diversity) and to base our study on a more general and consistent concept, “biodiversity characterizes qualitative, quantitative and functional aspects of biotic units at various levels of organization in a concrete or abstract context, and at a given temporal or/and spatial scale” (Beierkuhnlein, 2003). In consequence, species richness and metrics that are based on it are important, but they represent just one aspect of biodiversity. In fact, the total number of species co-occurring in a given community (*α*-diversity) is nested within the total number of a species pools occurring for instance at the landscape level (*γ*-diversity). But the reduction of biodiversity to the perspective of inventory and proportion would not cover spatial gradients in composition and species turnover (differentiation, *β*-diversity) (Jurasinski et al., 2009; Baselga, 2012) and also ignores functional diversity (e.g. functional traits), which is the main driver of ecosystem functioning.

In general, *β*-diversity is a crucial measure, since, given the same local richness of different sites (*α*-diversity), it directly considers the turnover among them. As an example, let A and B be two sampling sites with 10 different species each. If all 10 species are fully shared, the total *γ*-diversity would equal 10 species, while if all 10 species are completely different from one site to the other (high turnover, high *β*-diversity) the total diversity of the whole area based on the two focal sites would double.

Therefore, it becomes particularly interesting to understand how *β*-diversity originates, investigating how species composition differs among sites. In fact, species composition could be related to environmental conditions, or it could randomly fluctuate. A generally accepted hypothesis suggests that *β*-diversity might change as a function of species types living in a certain community. For instance, *β*-diversity should be small when communities are dominated by a limited number of competitive species; this is recognized as the null hypothesis and it entails a uniform distribution in species composition (Legendre et al., 2005).

The *β*-diversity concept reflects the environmental heterogeneity between sites and thus within a given larger area that contains several of the focal study sites. Heterogeneity is in fact highly associated with a high degree of biological diversity since heterogeneous sites offer a diversity of ecological niches (sensu Elton (1958)) that can be occupied if the species pool offers the respective ecological diversity to address these niches (Gaston, 2000; Rocchini et al., 2010). Furthermore, since *β*-diversity can be described as the spatial turnover among sites within a given region, it captures a fundamental feature of the spatial pattern of biodiversity.

In some cases, spatial turnover can result from local extinction processes that affect certain species more than others and enhance the dissimilarity between sites without dispersal (Steinitz et al., 2006). This is the case in highly fragmented landscapes where dispersal is limited (Hobbs et al., 2006). Even stochastic processes (sensu Moran (1950) and Clark (2008)) may enhance *β*-diversity in previously homogeneous ecosystems. For instance, sudden fragmentation (Alados et al., 2009) can lead to disfunctional source-sink metapopulations with intrinsic influences on the degree of spatial (and genetic) connectivity of organisms (Waples and Gaggiotti, 2006), resulting in the local loss of sink populations. However, in most situations, the spatial turnover and therefore the dispersal of species between sites (metapopulation and metacommunity dynamics) is linked to the distance among sites. Strictly speaking, the similarity between two sites decays with increasing distance between them (Rocchini, 2007), a process also known as the distance decay in similarity or the Tobler’s first law of geography (Tobler, 1970).

Hence, modelling the distribution of *β*-diversity in space is based on softening the role of individual species, which are not even completely described at wide geographical scales, for the sake of estimating a more efficient proxy for ecosystem patterns and processes. When these estimates are available from remote sensing data, this process can lead to rapid, large-scale monitoring and support management intervention aimed at preserving entire ecosystems, as stipulated by the Aichi Biodiversity Targets (https://www.cbd.int/sp/).

Field-based studies require an enormous investment in time in order to collect reliable biodiversity data. A pioneering example is the public database of the Global Biodiversity Information Facility (GBIF, https://www.gbif.org/). GBIF is a network funded by the world’s governments which contains almost 41,000 databases of species occurrences spread out across the world. The large amount of publically accessible data and the available techniques to analyze them will certainly facilitate biodiversity assessment for the areas that it covers. Unfortunately, however, although it would be possible in principle to use these data to make reasonable assumptions about biodiversity over larger areas, there are several limitations due to their quality (Maldonado et al., 2015). The errors that usually arise from field data are due to: (i) lack of or erroneous geographic coordinates of the sampling sites; (ii) incorrect taxonomic identification with poor quality control; and (iii) difficulties in proving a reliable random sampling with large areas being poorly covered. Furthermore, these data often appear as point data, while grids are usually used in order to synthesize diversity metrics. In addition, these data are mainly collected from presence-only data without any link to relative abundance, dominance, biomass or cover, which instead, is reflected in remote sensing. Finally, GBIF data are inadequate for local estimates of biodiversity as they do not consider co-occurrence data. Indeed, and contrary to recent databases at the community level such as the European Vegetation Archive (EVA) (Chytry et al., 2016) or the sPlot initiative (Bruelheide et al., 2019), GBIF does not provide information on species co-occurrence which is very problematic for biodiversity assessment and monitoring. Despite the disadvantages that come from the use of public databases, there is some benefits in the use of such data. First of all, there is a huge amount of data collected and provided by citizens and research institutions available in the GBIF database when compared to the data that could be collected locally, resulting in a huge saving of time and costs. More-over, GBIF data are standardized to the same format and therefore ready to use.

To overcome the issues due to the collection and availability of in situ ecological data, remote sensing imagery has become more and more important and is now considered a reliable tool to assess and monitor biodiversity (Tuanmu and Jetz, 2015).

### 1.2 The spectral species concept

Remote sensing based approaches have proven to be useful modelling techniques to detect the variability of biodiversity in space and time across scales of biological organisation, at different grains (spatial resolutions) and extents (Rocchini et al., 2013). Airborne sensors have even been used to detect and map single species distributions (Skorownek et al., 2017a), even the most tiny and inconspicuous ones such as *Campylopus introflexus*, a moss species which is highly invasive in Europe (Skorownek et al., 2017b).

Remote sensing techniques have been used to study the impact of land-scape and environment on biodiversity, and to explore and visualize spatial data and biodiversity change. Therefore, remote sensing data have become among the most time and cost effective tools, allowing to make relevant conservation actions in a relatively short period of time. Furthermore, remote sensing demonstrated the impact of biodiversity (including non-native invasive species) on ecosystem functioning (Ewald et al., 2018).

In general, vegetation absorbs the blue and the red light, for photosynthesis, while it reflects near infrared (hereafter, NIR) radiation due to the physical structure of the cells composing the leaf mesophyllum (Wegmann et al., 2016). The bands relative to RED and NIR are used as proxies for photosynthetic activity of the vegetation. These bands are usually incorporated in a widely used index, the normalized difference vegetation index (NDVI), which is calculated as NDVI=(NIR-RED)/(NIR+RED). The higher the relative abundance of photosynthetic vegetation, the higher would be the reflectance in the NIR band and the absorption in the RED band. NDVI ranges from −1 to 1, with 0 values usually associated with non vegetated areas and negative values associated with water surfaces or snow.

This index has widely been used to discriminate different vegetation types over an area. In fact, in several studies, NDVI is positively correlated to the net primary productivity (NPP, e.g. Gillespie et al. (2008)). Therefore, it can be used as a proxy to quantify species richness and diversity, based on the species-energy theory, proposed by Currie (1991), namely a relation between species richness and energy, that would depend mainly on annual potential evapotranspiration and actual evapotranspiration. Another hypothesis related to the variability in space of the spectral signal has been proposed by Palmer at al. (2002). The so called spectral variation hypothesis (SVH) states that the higher the environmental heterogeneity the higher would be the species diversity of an area, due to a higher number of available ecological niches.

Hence, based on the SVH, spectral variability can effectively be related to environmental heterogeneity and therefore it could be used to assess species biodiversity of an area. In this sense, since the spectral variability is derived from the information present in the pixels of an acquired image, it is important that the pixels, describing the area of study, would have a spatial resolution coherent with the ecological assumptions taken into account and such that predictions on biodiversity can be made.

Among the most novel methods to estimate diversity by remote sensing, described in Rocchini et al. (2018), the spectral species concept (Féret and Asner, 2014) is one of the most powerful, since it allows to couple k-means approaches to the gridded data obtained from remote sensing technologies as a mean to derive *α*- and *β*-diversity 2D-matrices. The spectral species algorithm allows the separation of the spectral space in subunits identified as spectral species. Its root theory is built upon two major founding principles. The first is the aforementioned Spectral Variation Hypothesis, relating spectral to environmental heterogeneity. The second is based on the plant optical types proposed by Ustin and Gamon (2010). This concept is mainly related to the use of particular sensors providing high spatial resolution images and able to measure different signals about the phenology, the biochemistry and the structure of vegetation. Such sensors can obtain information at the individual plant scale level.

The method is based on an unsupervised clustering algorithm, first relying on dimensionality reduction obtained after running a principal component analysis (PCA) and then on the actual clustering of the pixels, with the subsequent assignment to spectral species, based on a k-means approach. PCA and similar clustering methods have already been shown to reliably reduce the multidimensional spectral sets for models on species and biodiversity distribution (Rocchini et al., 2010). Furthermore, the method provides an interesting visual inspection of diversity building *α*- and *β*-diversity maps.

As far as we know, the spectral species concept has been applied so far only at the local scale (Féret and Asner, 2014). Hence, the aim of this manuscript is to extend this concept over wider spatial extents passing to a spectral community concept, by generating a heterogeneity map at a wide geographical scale to estimate *α*- and *β*-diversity across Europe.

## 2 The algorithm

The spectral species algorithm was originally developed to map tropical forest canopy diversity using imaging spectroscopy with a spatial resolution up to 2 meters (Féret and Asner (2014), Figure 1). Following the hypothesis that species are spectrally separable (Asner and Martin, 2009), the approach is based on the segmentation of the spectral space defined by the remote sensing data. In fact the spectral space is assumed to be a combination of several subspaces, reflecting the “signature” of one or several species. Therefore these subspaces would be the expression of a more general “spectral species”. From the resultant “spectral community”, it would be possible to derive the diversity of an area. Therefore, the output of this algorithm is not a list of the actual species of an area, but rather a map of the distribution of the spectral communities available within the area that may be used to calculate several diversity indices. In particular we focused here on *α*- and *β*-diversity metrics. Both introduced by Whittaker (1972), the first reflects the mean species diversity in sites at a local scale whereas the second is an indicator of the spatial (or temporal) heterogeneity at a relatively larger scale. In the algorithm, *α*-diversity is calculated in a neighbourhood (plot) of *n* × *n* pixels by the Shannon diversity index (Shannon, 1948) calculated as follow:

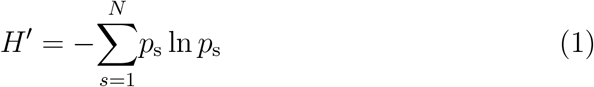

where *p*_s_ is the proportion of each spectral species *s* in each plot.

The *β*-diversity indicator is instead computed by the Bray-Curtis (here-after BC) dissimilarity metric (Bray and Curtis, 1957):

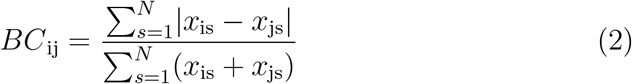

where *BC*_ij_ is the dissimilarity between plots *i* and *j* and *x*_is_ and *x*_js_ are the abundances of spectral species *s* in plots *i* and *j*.

**Figure 1:**
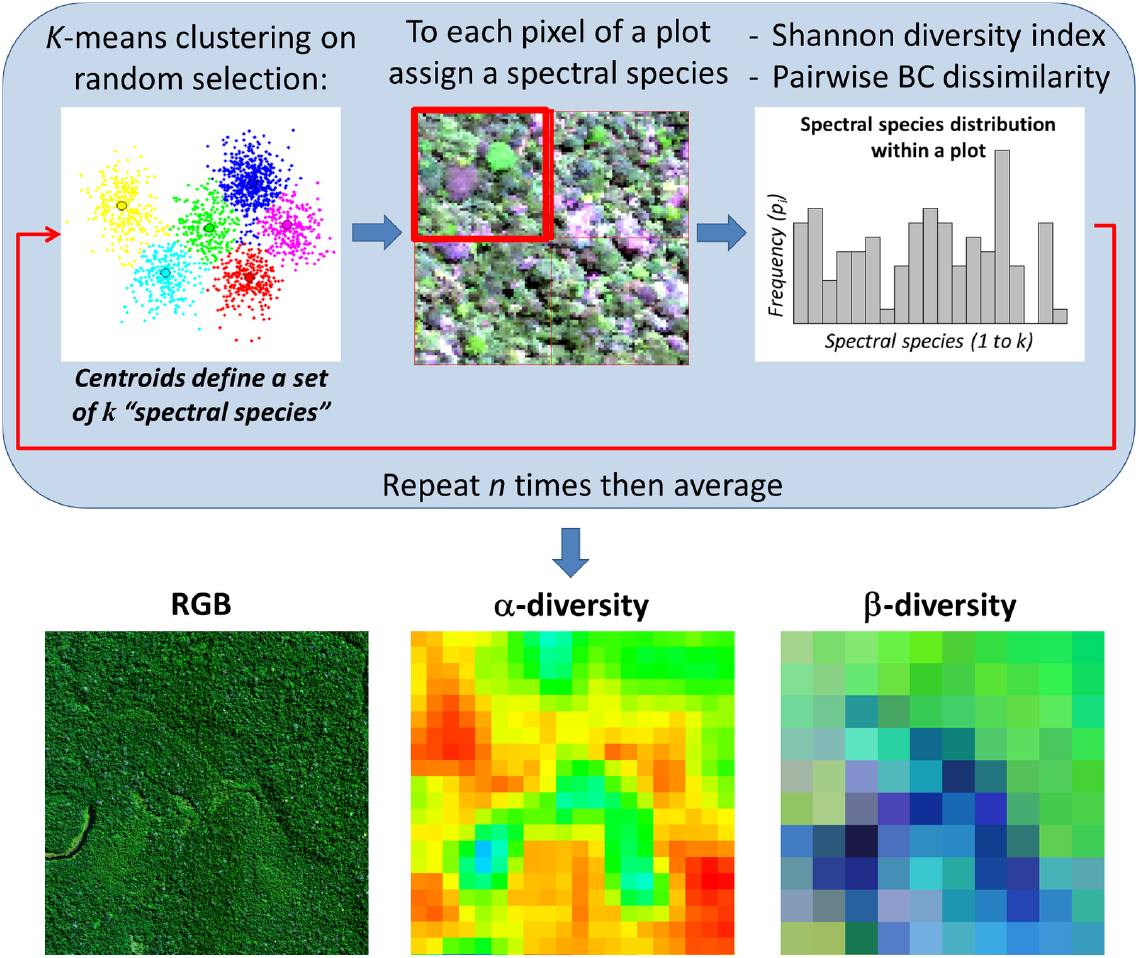
Diagrammatic representation of the steps of the algorithm used to achieve *α*- and *β*-diversities, redrawn from (Féret and Asner, 2014). Pixels are clumped in a spectral species and spectral community diversity is calculated. We refer to the main text and to Box 1 for additional information.

In the spectral species algorithm, once the BC dissimilarity matrix between all pairs of plots is computed, a multidimensional scaling is performed in order to translate information about the pairwise dissimilarity among P plots into a configuration of P points mapped in a 3-dimensional Cartesian space such as NMDS or PCoA (Mead, 1992). This simplified translation of the BC dissimilarity matrix can then be displayed as a colored map. More details can be found in Féret and de Boissieu (in press).

While the Shannon index has a theoretical maximum limit corresponding to the *ln*(*richness*), the Bray-Curtis index ranges from 0 to 1, where 0 is indicating that the two sites are identical whereas 1 indicates that the two sites do not share species. Hence, BC can be considered as an estimate of the heterogeneity of a certain area. The final aim of the method was to generate an heterogeneity map across the study region. Strictly speaking, the method is a clustering approach which (i) divides the subspaces in spectral units and (ii) assigns it to spectral species from which (iii) different diversity maps can be obtained. Box 1 focuses in detail on the main steps of the algorithm, while the dedicated R package biodivMapR is now available (https://github.com/jbferet/biodivMapR) and fully described in Féret and de Boissieu (in press).

### 2.1 Application of the algorithm

Remote sensing data are usually provided as raster objects with a geographic coordinate system information, namely regular grids (matrices) or stacks of raster layers (e.g. one raster layer per band for multispectral or hyperspectral data), in which each cell represents a pixel with the corresponding reflectance value associated to a specific band. Such data have been manipulated with the Software R Development Core Team (2019). R can be used for remote sensing data analysis since it includes spatial functionalities throughout a suite of R packages like the `rgdal` and `raster` packages (see Box 2 for more information).

Our main purpose was to apply the spectral species algorithm to a continentalscale geographical region such as Europe. Hence, Moderate Resolution Imaging Spectroradiometer (MODIS) data, with a spatial resolution of 500m covering Europe, were downloaded from the United States Geological Survey (USGS) site (https://lpdaac.usgs.gov/products/mod09a1v006/). After a visual check of the images, in order to guarantee i) the coverage of a complete phenological period and to ii) avoid noise related to clouds, we referred to the RED and NIR bands from 2018 from January to December (one image per month), to calculate NDVI, by generating a sample set of 12 NDVI images (Figure 2).

**Figure 2:**
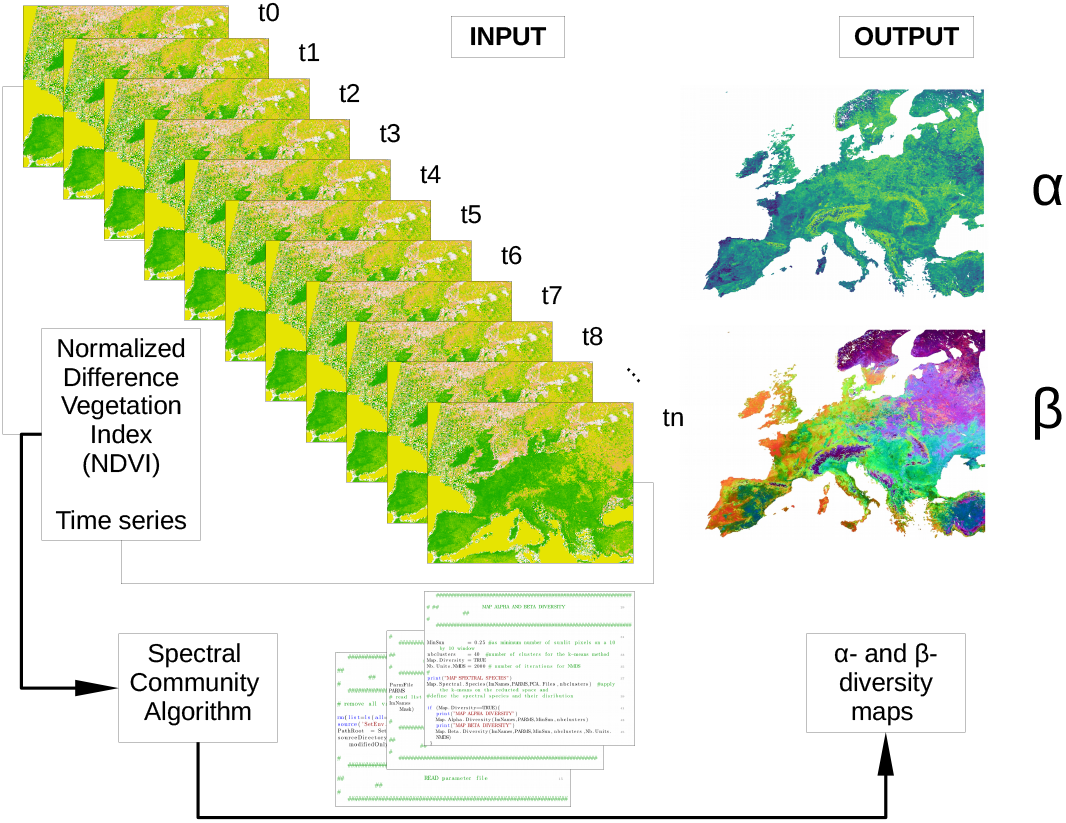
An input set of *n* images can be handled to create a time series and use the stack to further calculate the spectral community diversity. In our paper, a stack of 12 NDVI images of 2018 from the MODIS sensor was processed by the spectral species algorithm, by producing *α*- and *β*-diversity maps.

Due to the coarse spatial resolution of MODIS images (500m), the reflectance related to a single plant species is averaged and mixed with the reflectance of other species within a single pixel. In other words, the direct relationship between spectral species detected in the spectral space versus the number of plant species does not hold true. However, in any case, from a diversity measurement perspective, this is just a matter of terms being used, with spectral species being actually more related to field plant communities, habitats or other ecological entities.

For the derivation of spectral species, in order to define the number of clusters, we relied on the highest number of clusters with stable results after a trial and error procedure, reaching 200 clusters, i.e. spectral species. Once pixels with similar NDVI values in 12 dimensions were clumped together, Shannon’s *H*′ was calculated with a window size of 10×10 pixels and an output resolution of 5km. The attained *α*-diversity map quantitatively showed the local spectral diversity distribution over Europe (Figure 3), with a higher heterogeneity found in i) more topographically complex regions, mainly due to strong local differences induced by elevation gradients (passing from forests to grasslands, to rocks and snow), and/or differences in terms of seasonality in relation with elevation, as in Rocchini et al. (2019), and in ii) more contrasted agricultural areas in both the spatial and temporal dimensions (Hobbs et al., 2006; Vihervaara et al., 2017). Concerning topographical complexity, the higher variability in areas with a marked topographical gradient might be related to shadows. Local field work in such areas will be needed to validate the measurements in such areas.

**Figure 3:**
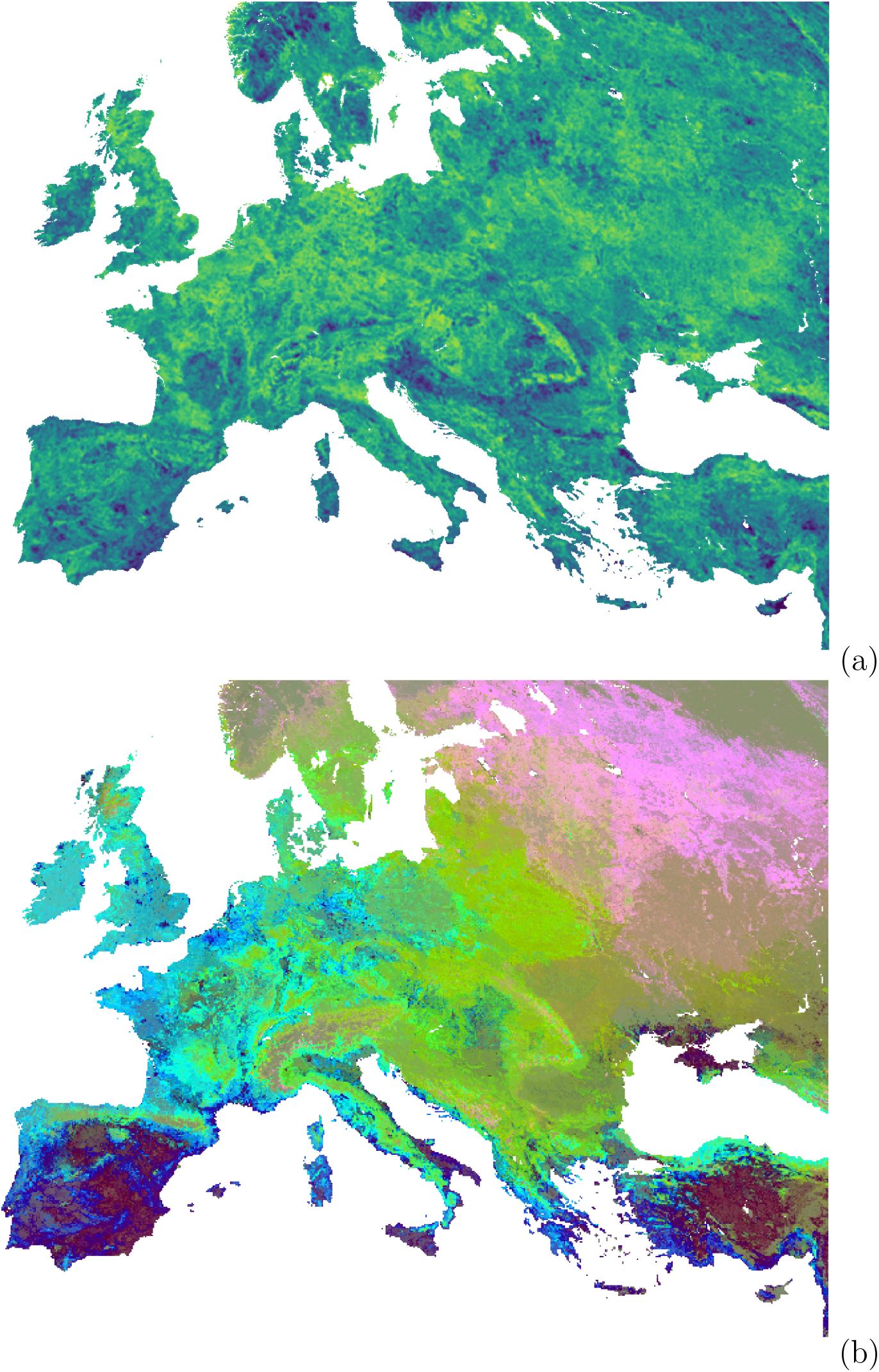
*α*- (a) and *β*-diversity (b) maps obtained by the spectral species algorithm. (a) The *α*-diversity map, based on Shannon’s *H*′ index (ranging from blue [low values] to light green [high values]) calculated in a 10×10 pixels local neighbourhood, corresponds to the local entropy of clusters, so that each location is independent from the others; (b) The *β*-diversity map - Bray-Curtis dissimilarity reduced to 3 dimensions with NMDS - provides information about the dissimilarity a2m6ong any location in the image. Here, the distance between pairs of spatial units is expressed as a 3 colour code.

*β*-diversity (Figure 3) showed a clear differentiation among different areas over Europe. The attained map was in line with the European Environmental Agency (EEA) map of ecoregions (Figure 4, see Mucher et al. (2009)). The correspondence of the achieved patterns in the two maps was apparent, with a similar contour of the major ecoregions such as the mediterranean, the atlantic, the continental, the boreal and the alpin regions. This demonstrates an intrinsic ability of the spectral species approach to capture differences in the physiological and functional properties of vegetation even at wide spatial scales, starting from spectral reflectance or spectral indices. Minor differences were mainly related to the biogegraphical (i.e., purely spatial) differentiation of ecoregions in the EEA map. As an example, north and south alpine ecoregions could not be distinguished by the spectral species approach, since they both have very similar conifer species composition, with the same physiological, phenological and thus spectral pattern.

**Figure 4:**
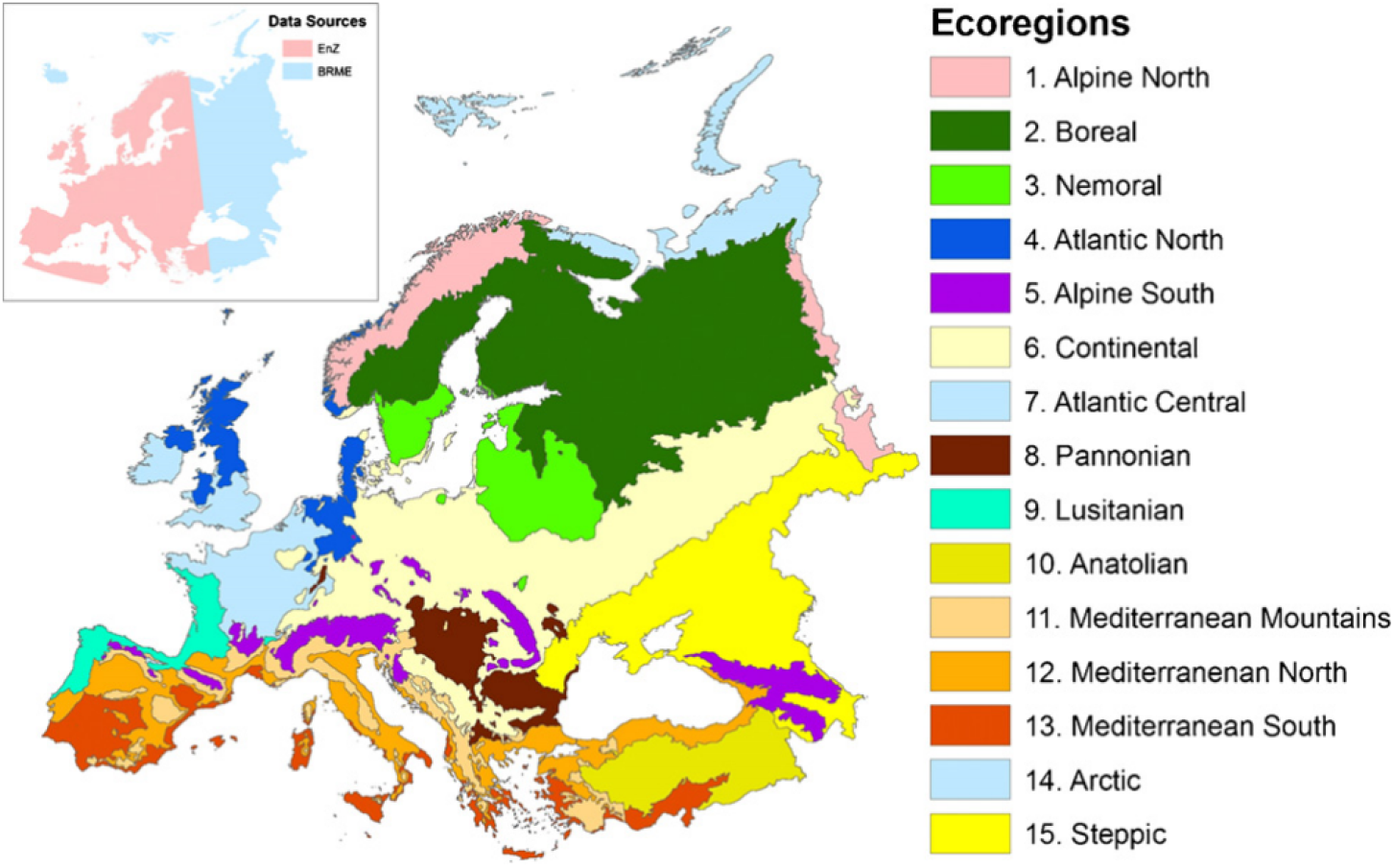
European Environmental Agency ecoregions map redrawn by Mucher et al. (2009). Similar maps at a coarser grain are provided by Mouchet et al. (2015) and Dinerstein et al. (2017).

## 3 Discussion

In this paper, for the first time, the spectral species concept has been extended from the consideration of a single species to an entire community. We demonstrated that the combined use of the novel unsupervised clustering method proposed by Féret and Asner (2014) with NDVI time series at European scale, allows the derivation of local (*α*) diversity and turnover (*β*) relying on free to use and operationally available satellite data.

With regards to a potential validation with in-situ data, the uncertainty of wide-scale datasets hampers a spatial overlap. In this case, in-situ datasets meet all five major concerns recently raised by Hobohm et al. (2019), i.e.: i) there is insufficient data coverage across Europe to make an unbiased comparison between predicted and actual distributions, ii) taxonomic standards differ across sampled regions, iii) there are generally different shapes of areas being sampled, iv) political borders often define sampling areas and aggregated sampling areas, and v) data are not aggregated in the same way in all areas. Furthermore, spatial information has an intrinsic varying degree of relevance mainly due to the fact that, rather than species lists, it is composed of geometrical precision, attributes robustness and temporal consistency (Hobona et al., 2006). Finally, different models and approaches to measuring diversity inevitably provide different outputs, as pointed out in the generalised entropy theory put forward by Rényi (1961). Given the above validation difficulties, we decided to qualitatively compare our generated output, in particular the *β*-diversity map, with existing ecoregion maps, which are expected to discriminate different spatial areas based on natural borders defined by biological diversity (https://ecoregions2017.appspot.com/) and thus are intrinsically related to differences in the species and spectral turnover of communities.

Since the output of the algorithm represents the variation of the pixel values in space and time, the most diverse pixels were those with the highest turnover among the neighborhood areas and most affected by seasonality. The importance of accounting for turnover instead of simple richness has been widely discussed in the ecological literature (Tuomisto, 2010), since environmental variability over spatial gradients is one of the major drivers of the structure and composition of diversity (Legendre et al., 2005). In this view, the use of the “spectral species concept”, defined as the variation of clustered pixel values, represents a powerful approach for the investigation of gradient variation of diversity in space and, potentially, in time.

In general, the measure of variability in space has been demonstrated to follow scale-based differentiation. In other words, results are expected to change with spatial scale in terms of both grain (spatial resolution) and extent (extent of geographical area of interest, Palmer at al. (2002)). Regarding extent, one of the major weaknesses of the proposed algorithm in *β*-diversity quantification (although this applies in general to all measurements of turnover) is that by increasing the extent of an observation area, the estimated values for an individual comparison between sites are modified by the increasing spectral species pool.

Additional drawbacks at the current stage of the algorithm include: i) the use of remotely sensed data which are not necessarily related to the main drivers of species distributions and of diversity, ii) the general multicollinearity found in most of the remotely sensed sets, iii) the unsupervised clustering process being adopted.

Concerning climate, a solution might be found in the use of remotely sensed derived climate data adding climate change as an additional layer of complexity as in Rocchini et al. (2015a) and in (Zellweger et al., 2019). Also in this case multicollinearity of climate variables should be seriously taken into account, as we did for the original remote sensing data, by applying a PCA to reduce the noise in the data and detect potential artifacts; consequently, PCA components might also be visualized to find potential congruence between spectral species and real species patterns. Finally, the process for grouping pixels in spectral species is based on an unsupervised clustering, where the definition of the number of clusters should be done a-priori. In this case, we hypothesized that the diversity of types of landscapes and gradient of climates across Europe may require a large number of clusters to correctly differentiate among them, relying on a fuzzy view of ecosystems (Rocchini and Ricotta, 2007). Hence, we decided to adopt a trial and error procedure until a threshold was reached in which no significant changes were found. Such a threshold resulted in 200 clusters. In the near future, it would be interesting to make a sensitivity analysis to demonstrate the impact of the number of clusters on the final analysis.

Considering the use of remote sensing for species diversity estimates, correlation and determination coefficients are generally statistically significant but low, hampering the direct use of remotely sensed diversity in simple univariate models (Rocchini et al., 2018). In fact, the relationship between *α*- or *β*-diversity and habitat heterogeneity, which is the founding principle for the use of remote sensing data for these analyses, is rarely linear (Ferrier et al., 2007), mainly because of variation in the rate of species turnover along an environmental gradient. However, remotely sensed variables are generally well suited in more complex multivariate models accounting for part of the diversity explained for species communities (Rocchini et al., 2018). This is especially true considering that environmental turnover generally explains more variation in species diversity rather than mere spatial structure (Hernandez-Stefanoni et al., 2012). Moreover, based on their high temporal resolution, remote sensing data might be useful to detect drastic changes of diversity in space and time, e.g. related to catastrophic events, overall considering the intrinsic difficulties in relying on in-situ data for wide geographical scales (Cord and Rödder, 2011; Hobohm et al., 2019).

From an ecological perspective, remote sensing imagery bands (dimensions) show a high affinity with the hypervolume axes proposed by Hutchinson (1957) for modelling species niches. In the Hutchinson’s theory, an hypervolume is represented by a space defined by a set of *n* independent axes which could be related to the final variables driving the realised niche of a species (see also Blonder (2017) and Ricotta et al. (2010) on the niche differentiation concept). In our case, such axes would be the original satellite sensor bands being strictly related to the identification of a spectral species and the resulting spectral community in a site, instead of a niche. From this point of view, spectral species and communities are in line with joint species distribution models (JSDMs), which explicitly take into account biotic interactions among species in a community, while in our model the “interaction” among pixel values is ruled out in general by their proximity both from a spatial and from a spectral point of view. In this paper, the final aim was not to model single spectral species or spectral communities but rather to estimate diversity and its change over space and time, following the mathematical principles described in Liu et al. (2014) and Rocchini et al. (2015b), for which the distribution of diversity over space is actually a particular case of the so-called switched systems, i.e. hybrid systems resulting from both continuous and discrete dynamics with a high number of different potential variables acting as main drivers of diversity response. In our view we succeeded here to fill a previous gap in spatio-ecological analysis, i.e. the translation of what in remote sensing science is known as “spectral mixture modeling” (Jensen, 2015) into an ecological diversity theory approach. In spectral mixture modeling the measured spectral reflectance is decomposed as a mixture of endmembers. In our case, such a mixture was used to directly compute alpha- and beta-diversity over wide spatial areas in a very short time.

## 4 Conclusion

Predicting and mapping *α*- and *β*-diversity using remotely sensed images acquired over large areas is currently a key topic in ecology, and could provide landscape managers with rapid and effective tools to estimate and monitor global change. In this paper, we proposed a novel method based on preliminary unsupervised clustering of spectral data (NDVI time series derived from MODIS data), assigning each pixel to a “spectral species” and then calculating diversity based on a dissimilarity metric. At the scale of this study, the one-to-one relationship between spectral species and in-situ plant species is not achieved, but the spectral species concept still holds true once considering that the detected spectral species in the spectral space are related to higher-order plant hierarchies (assemblages, entire habitats, etc.). That is, from an algorithmic point of view, the bulk of the calculations are unaltered. Based on the results presented here, the use of the spectral species and communities concept would appear to promote more effective planning and policies related to the conservation of wild species, by improving our understanding of the dynamics of local and global biodiversity at various spatial and temporal scales.

## 5 Acknowledgements

DR was partially supported by the H2020 project SHOWCASE and by the H2020 COST Action CA17134 “Optical synergies for spatiotemporal sensing of scalable ecophysiological traits (SENSECO)”. This paper has been partially developed during a Short Term Scientific Mission supported by the H2020 COST Action CA15212 “Citizen Science to promote creativity, scientific literacy, and innovation throughout Europe”.

JBF and FdB acknowledge financial support from Agence Nationale de la Recherche (BioCop project—ANR-17-CE32-0001) and TOSCA program grant of the French Space Agency (CNES) (HyperTropik/HyperBIO project).

## Box 1 - Steps composing the spectral species algorithm

1. A Principal Component Analysis (PCA) is applied to the spectral data. PCA is not performed on the whole image, but only on a large subset of pixels randomly selected from the image. Due to the high dimensionality of the data, the reduction of the dataset is not altering the result. Those principal components explaining most of the variance of the original set are then retained for further steps.
2. A subset of pixels is then randomly selected across the entire map and the spectral space containing such a subset is partitioned into spectral species using k-means clustering with the number of k clusters being decided a priori. Then the centroids defining the spectral species are located.
3. The spectral dataset is divided into final mapping units. Each pixel is assigned to a given spectral species based on the minimal Euclidean distance between pixels (Peuquet, 1992) and the previously defined centroids.
4. A spectral species distribution is obtained for each mapping unit from which the *α*- and *β*-diversity indices are computed as previously stated.
5. Since the spectral species distribution is obtained by a subset of pixels, in order to avoid under-representation of some small-scaled ecological classes (e.g. small scale vegetation patterns), steps 4 and 5 are repeated 100 times, and the indicators obtained for each repetition are averaged. In particular the Bray-Curtis dissimilarity matrix is computed for each pair of spatial units, based on their spectral species distribution at each iteration; then the final matrix corresponds to the BC dissimilarity averaged over all the iterations.
6. Non metric Multidimensional Scaling (NMDS) (e.g. Borg and Groenen (2005)) is applied to the matrices in order to obtain a visual representation of the results. NMDS is an ordination technique usually applied in ecology that differs from other ordination techniques as PCA, since in NMDS a small number of axes are chosen prior to the analysis and then the data are fitted into the chosen dimensions. Furthermore, NMDS is not an analytical but numerical technique, seeking for the right solution (convergence) iteratively. Finally, NMDS is not an eigenvector-eigenvalue technique, hence a NMDS ordination can be rotated among the axes. NMDS is mostly used in ecology for its versatility since it accepts any distance measure of the samples. In this case the Bray-Curtis matrix was used. In the applied NMDS approach, the first step is generally to decide the number of reduced dimensions; in this case 3 dimensions were chosen. The algorithm starts with the construction of initial random arrangements of the pixels. Then the Euclidean distances among the samples is calculated in this first configuration; those distances are regressed against the original distance matrix, and the predicted ordination distances are calculated. Finally, the regression is fitted by the least-squares method. The goodness of fit is measured by the sum of squared differences between ordination-based distances and the predicted distances. The goodness of fit is calculated through the Kruskal’s Stress index:

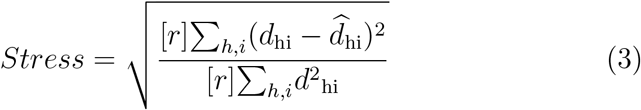

where *d*_*hi*_ is the ordinated distance between pixels *h* and *i*, and 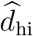 is the distance predicted from the regression. Then, a new configuration is computed moving in the direction in which stress changes most rapidly. The entire procedure is repeated until convergence. A *Stress* value that provides an excellent representation in the reduced dimensions is considered to be lower than 0.05; nevertheless a value of *Stress* < 0.2 is still considered a good representation Borg and Groenen (2005).

Basically, the algorithm provides both single spectral species maps and the *α*- and *β*-diversity maps. The algorithm input file needs to be in ENVI binary format with the corresponding header file. The file should be in Band Interleave by Line (BIL) format and 2-byte signed integer, and should not have extension. A further masking file in the same format is necessary in order to mask clouds and water surfaces.

## Box 2 - Packages used in this manuscript to handle and analyse spatial data in R

- `raster`: It provides classes and functions to manipulate geographic data in raster format. Raster data divides space into cells (as pixels) of equal size (in units of the coordinate reference system). Along with the `raster` package, the `sp` package is also loaded, which provides spatial object classes and methods to retrieve coordinates.
- `rgdal`: It provides functions to import ad export spatial data in different formats.
- `RStoolbox`: A toolbox for remote sensing image processing and analysis.
- `rasterdiv`: It provides algorithms for measuring diversity from spatial matrices.

